# Lactate:propionate molar ratio and substrate concentrations shape valerate production in secondary lactate fermentations

**DOI:** 10.64898/2026.05.06.722830

**Authors:** Ángel Estévez, Ramon Ganigué

## Abstract

Odd-chain carboxylates such as valerate and heptanoate are ecologically relevant metabolites and promising platform chemicals, yet the factors leading to their formation during secondary lactate fermentations remain poorly understood. Here, a continuous anaerobic bioreactor was operated for 297 days at pH 5.5 to evaluate how lactate:propionate molar ratios shape product spectrum in lactate fermentations. Valerate was the predominant odd-chain product under all conditions, reaching concentrations up to 110 mM, while heptanoate accumulated only at low levels (<10 mM). At low lactate concentrations (110-220 mM), product selectivity strongly depended on the lactate:propionate ratio. When lactate:propionate ratios were around 1.2 mol/mol, odd-chain products were favored, whereas higher ratios (up to 4.8 mol/mol) shifted metabolism toward caproate and butyrate formation. However, this trend was not maintained at higher lactate concentrations (330-440 mM; lactate not fully consumed), where odd-chain selectivities remained high even at lactate:propionate ratios of 4.8 mol/mol. Pathway analysis allowed to infer that under high-lactate conditions, up to 30% of lactate was used for propionate and acetate formation. Microbial community analysis revealed a stable dominance of *Caproiciproducens* spp. when valerate was the main product. Overall, this work provides mechanistic insights into the ecology of lactate fermentations and offers a framework for steering product selectivity in engineered anaerobic systems.

**Highlights:**

- Valerate was the dominant product in most tested conditions, reaching up to 110 mM.
- Lactate:propionate ratio and substrate concentrations drive product selectivities.
- High lactate concentrations were associated with inferred endogenous propionate formation.
- *Caproiciproducens* dominance was associated with sustained valerate production.

## 1. Introduction

Microbial chain elongation is a widespread process observed in natural environments such as soils, wetlands, and the gastrointestinal tract^1^. More recently, it has also been proposed as a technology for upgrading waste and other low-value feedstocks into valuable products^2,3^. Microbial chain elongation typically relies on the β-oxidation pathway, in which short-chain carboxylates are elongated by two-carbon units per cycle, typically using ethanol, lactate, or sugars as electron donors^4^. Among these, lactate is of particular interest because of its role as a central intermediate in many anaerobic ecosystems^5,6^. It is an abundant intermediate of carbohydrate fermentations by lactic acid bacteria and bifidobacteria, and it rarely accumulates in natural environments for extended periods^7^. Instead, lactate is rapidly consumed by secondary fermenters, and is either elongated into carboxylates such as butyrate, valerate, and caproate or fermented into acetate and propionate via the acrylate or methylmalonyl-CoA pathways^8^. Previous work has shown that environmental conditions can steer lactate fermentation between chain elongation and propionate production. For example, pH was identified as a key factor governing the competition between lactate-based chain elongators and propionate producers^9^. At pHs below 6, known chain elongators like *Caproiciproducens* spp. were enriched, leading to the production of elongated carboxylates such as caproate. In contrast, more neutral pH values favoured the selection of *Veillonella* spp. and shifted product formation towards propionate and acetate.

To date, most lactate-based chain elongation studies have focused on the production of even-chain carboxylates, e.g. caproate and caprylate^10–16^, while the production of odd-chain carboxylates remains comparatively poorly understood^17,18^. Odd-chain products such as valerate and heptanoate are nevertheless of broad relevance, both because of their potential as key intermediate chemicals (e.g. PHBV production^19^) and because of their ecological occurrence. Valerate is frequently detected in gut ecosystems, where together with propionate they represent the dominant odd-chain carboxylates. Valerate has been shown to positively influence host physiology^20^, but only recently attention has also been placed on how this valerate is produced in the gut^21,22^. In natural fermentations and mixed-culture bioprocesses, valerate is also regularly observed even when propionate is absent in the substrate, suggesting that pH alone may not fully explain the balance between lactate-based chain elongation and propionate formation in mixed communities. For instance, Kucek et al. (2016) and Candry et al. (2020) still observed significant odd-chain carboxylate production at acidic pHs in continuously fed bioreactors with lactate as the only carbon and energy source^9,14^. Similarly, Allegue et al., (2022) reported up to 20% of valerate in the product spectrum of a carbohydrate fermenting CSTR at pH 5.5, which increased to 34% upon additional lactate supply^17^. Heptanoate formation, in contrast, is comparatively rare, with only few reports available, mostly involving ethanol as electron donor^23^.

The ratio between electron donor and acceptor has previously been shown to influence product selectivity in chain elongation systems, with higher lactate:acetate ratios leading to higher caproate production²⁵. However, the interpretation of the lactate:acetate ratio is complicated by the fact that acetate can be both supplied externally and generated from lactate oxidation, making it difficult to distinguish how much lactate flux is directed towards acetate production versus chain elongation. Evidence from ethanol-based chain elongation systems provides a clearer framework for understanding how the ratio between electron donor and acceptor can affect odd-chain elongation²⁴. Candry and co-workers demonstrated that the ethanol:propionate ratio strongly influenced product selectivity, with higher ethanol availability lowering the fraction of odd-chain products in both *Clostridium kluyveri* and mixed communities due to increased ethanol oxidation to acetate. This is explained by the fact that in reverse β-oxidation, acetyl-CoA (formed from ethanol) is combined with a propionyl-CoA molecule during odd-chain elongation to eventually yield valerate. However, for every 5 molecules of acetyl-CoA used to produce valerate, one is further oxidised to acetate to generate energy for the cell. This acetate is then also available to be elongated to butyrate and caproate. Consequently, the higher the ethanol compared to propionate, the higher the fraction of even-chain carboxylates. Lactate can similarly provide acetyl-CoA for chain elongation through its oxidation to pyruvate and subsequent formation of acetyl-CoA. However, unlike ethanol, lactate can alternatively be converted to a mixture of propionate and acetate, making the interpretation less straightforward. Based on the stoichiometry described above, we therefore hypothesise that increasing the lactate:propionate ratio will also lead to a higher fraction of even-chain carboxylates.

The aim of this study was to determine how lactate:propionate molar ratios shape the product spectrum in secondary lactate fermentations. Overall, this work provides new insights into the ecology of lactate fermentations and valerate production and establishes a framework for selectively directing carbon flux towards odd-chain carboxylates in engineered anaerobic environments.

## 2. Materials and methods

### 2.1. Reactor operation

A lab-scale 1 L double-jacketed glass bioreactor with a working volume of 0.5 L was operated for 297 d as a CSTR with a hydraulic retention time of 4 d. The reactor temperature was maintained at 30 ± 1°C and the pH was controlled to 5.5 ± 0.1 with 1M HCl solution. Agitation was performed with a magnetic stirrer at a stirring rate of 400 rpm. A pH probe and pH control pumps were connected and controlled by an in-house made transmitter box. The reactor was inoculated with 50 mL of sludge from an anaerobic digester. After inoculation, the reactor was sparged with nitrogen gas to ensure anaerobic conditions. The reactor operation was monitored by online measurements (pH and temperature) and offline liquid and solid analyses. Suspended solids were separated from the mixed liquor by centrifugation (7317 rcf for 20 minutes). The supernatant after membrane filtration (0.2 μm pore size filters) was stored at −20°C pending liquid analyses.

### 2.2. Media composition

Lactate and propionate ratios and concentrations were varied along the experiment, as depicted in Table 1. The reactor was initially operated at a lactate:propionate ratio of 2.4 mol/mol. The ratio was then adjusted as shown in Table 1 while keeping propionate constant at 90 mM. The same lactate ratios were subsequently tested again by fixing lactate at 220 mM and varying the propionate concentration. Additionally, an extra test was performed at a molar ratio of 4.8 mol/mol, with 330 mM of lactate and 70 mM of propionate. All conditions were tested in a single reactor, so phase to phase comparisons should be interpreted in the context of potential reactor adaptation and changing substrate conditions. The growth medium contained the following compounds (in g/L): MgCl_2_·6H_2_O, 1; CaCl_2_·2H_2_O, 0.5; NaH_2_PO_4_·2H_2_O, 0.5; Na_2_SO_4_, 0.1; KCl, 1; NH_4_Cl, 2; yeast extract, 1; tryptone, 4; SL10 trace elements solution, 1 mL/L; Se-W trace element solution, 1 mL/L; 7-vitamins solution, 0.1 mL/L. The Se-W trace element solution consisted of (in mg/L): Na_2_SeO_3_·5H_2_O, 3; Na_2_WO_4_·2H_2_O, 4; while the 7-vitamins stock solution was as follows (in g/L): vitamin B12, 1; p-aminobenzoic acid; 0.8; D(+)-Biotin, 0.2 g; nicotinic acid, 2; Ca-pantothenate, 1; pyridoxine hydrochloride, 3; thiamine-HCl·2H_2_O; 2.

**Table 1.**
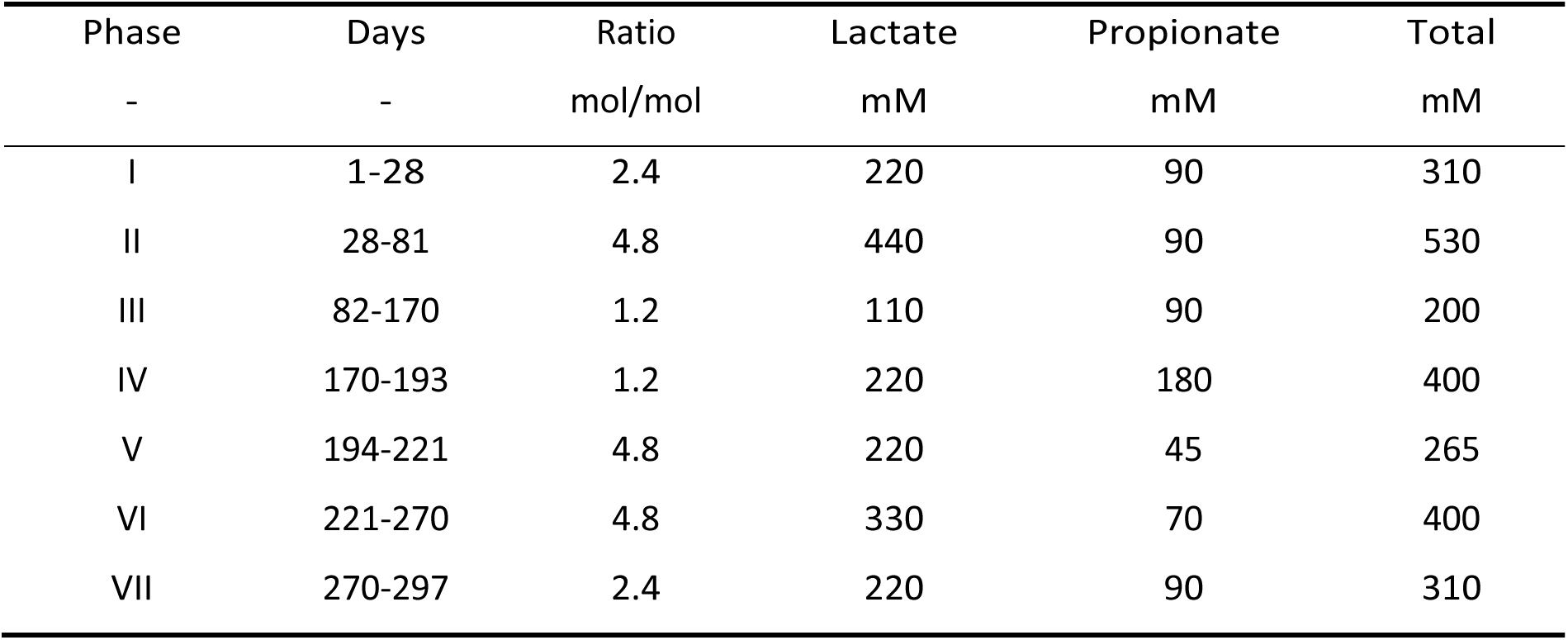
Overview of the lactate and propionate concentrations and ratios tested.

| Phase | Days | Ratio | Lactate | Propionate | Total |
| --- | --- | --- | --- | --- | --- |
| - | - | mol/mol | mM | mM | mM |
| I | 1-28 | 2.4 | 220 | 90 | 310 |
| II | 28-81 | 4.8 | 440 | 90 | 530 |
| III | 82-170 | 1.2 | 110 | 90 | 200 |
| IV | 170-193 | 1.2 | 220 | 180 | 400 |
| V | 194-221 | 4.8 | 220 | 45 | 265 |
| VI | 221-270 | 4.8 | 330 | 70 | 400 |
| VII | 270-297 | 2.4 | 220 | 90 | 310 |

### 2.3. Analytical methods

Lactate and acetate concentrations were determined by ion chromatography. Samples were analyzed with a 930 Compact IC Flex (Metrohm, Herisau, Switzerland) equipped with a Metrosep Organic acids 250/7.8 column, a Metrosep organic acids Guard/4.6 guard column and with a Metrohm CO_2_ Suppression and an 850 IC conductivity detector. Anions were eluted at a flow rate of 0.5 mL min^-1^ using 1 mM sulfuric acid. Carboxylic acids (C3 to C8, with their respective isomers) were analyzed by gas chromatography. Samples were analyzed with a GC-2014 (Shimadzu®, The Netherlands) equipped with a DB-FFAP 123-3232 column (30m/0.32mm/0.25μm; Agilent, Belgium) and a flame ionization detector. The carrier gas was nitrogen at a flow rate of 2.49 mL/min. Samples were conditioned with 2 mL sulfuric acid, 200 mg sodium chloride and methyl hexanoic acid was added as an internal standard before further extraction with diethyl ether (1:1 volume sample/ether). The sample (1 µL) was injected at 250°C with a split ratio of 50 and a purge flow of 3 mL min^−1^. The oven temperature increased by 10°C min^−1^ from 110 to 250°C where it was maintained for 5 min. The FID had a temperature of 300°C. Composition of produced gases was measured by gas chromatography using a Compact GC4.0 Global Analyzer Solutions, The Netherlands), equipped with a Molsieve 5A pre-column and Porabond column (CH_4_, O_2_, H_2_ and N_2_), and a Rt-Q-bond pre-column and column (CO_2_). Concentrations of gases were determined by means of a thermal conductivity detector. The carrier gases were N_2_ and He. The harvested biomass pellet dry weight and ash contents were estimated based on standard methods (105°C drying and 550°C ashing) and referenced to the sample volume for total and volatile suspended solids (TSS and VSS), respectively.

### 2.4. Carbon and electron balances and pathway analysis

The reactor was assumed to be in steady state when the product spectrum and the acid addition per day were constant (<10% variation) for at least 3 HRTs (≍12 days). Carbon and electron balances were set up to assess the accuracy of the measurements, and were evaluated by comparison of the consumed lactate and propionate with the fermentation products in the effluent, the off-gas and biomass formation. The potential contribution of yeast extract and tryptone was assessed by comparing the unidentified carbon and electron fractions in the influent and effluent. As these fractions remained similar, their net contribution to the carbon and electron balances was considered negligible. Biomass concentrations were determined from VSS measurements and were assumed to be represented as CH_1.8_O_0.5_N_0.2_^24^. The gap between the influent and the sum of the identified products was classified as “unidentified fermentation products”. Product yields were defined as the mol of formed product divided by the net consumed lactate and propionate. The estimated endogenous propionate production was not considered in the calculation of product yields. The contribution of endogenous propionate production in the reactors was assessed by comparing the moles of odd-chain products (valerate and heptanoate) with the moles of consumed propionate, as described in Equation 1, where LA, PA, VA, HA stand for lactate, propionate, valerate and heptanoate, respectively. For all calculations, the linear and branched isomers of each carboxylic acid were lumped together. These calculations further assumed that unidentified products and biomass incorporation did not substantially distort the balance, which appears reasonable given the reported carbon and electron balances.

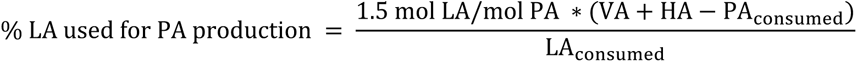

The endogenous propionate production was considered active when the mols of odd-chain products exceeded the consumed propionate, indicating that additional propionate must have been formed endogenously to support the production of valerate and heptanoate. The lactate required to drive propionate production was subsequently estimated based on its stoichiometry, in which three moles of lactate are converted into two moles of propionate, one mole of acetate, and one mole of carbon dioxide ^7^.

### 2.5. Microbial community structure

Fresh samples were collected from the reactor at selected time points during steady state conditions. The suspended solids were separated from the mixed liquor by centrifugation (2 min, 20238 rcf) and stored at –20°C until DNA extraction. DNA was extracted from 0.5 mL of the mixed sample using a FastDNA® SPIN kit for soil (MPBio, USA) according to the manufacturer’s instructions. The bead-beating step was performed using a PowerLyzer 24 Homogenizer (Qiagen, Hilden, Germany) at 2000 rpm for 5 min. Extracted DNA (10 µl) was sent to LGC genomics GmbH (Berlin, Germany) for library preparation and 16S rRNA gene amplicon sequencing on an Illumina MiSeq platform with v3 chemistry with primers 341F (5’-CCT ACG GGN GGC WGC AG −3’) and 785Rmod (5’-GAC TAC HVG GGT ATC TAA KCC-3’)^25^.

Illumina sequencing data was analyzed using R version 4.4.2 (2024-10-31). The DADA2 R package was used to process the amplicon sequence data according to the pipeline tutorial^26^. In a first quality control step, the primer sequences were removed, and reads were truncated based on a quality score threshold (truncQ=2). Besides trimming, additional filtering steps were conducted to exclude reads containing any ambiguous base calls or high expected errors (maxEE=2.2). After dereplication, unique reads were further denoised using the Divisive Amplicon Denoising Algorithm (DADA) error estimation algorithm and the selfConsist sample inference algorithm (with the option pooling =TRUE). The resulting error rates were inspected, and after approval, the denoised reads were merged. Subsequently, the Amplicon Sequence Variant (ASV) table obtained after chimera removal was used for taxonomy assignment employing the Naive Bayesian Classifier and the DADA2 formatted Silva v138.1 database^27^. In addition, any ASV with a total abundance across all samples less than or equal to 1 was excluded. ASV relative abundances were calculated by total-sum scaling, by dividing the read count of each ASV by the total number of reads in the corresponding sample. No rarefaction was performed. The raw fastq files of the 16S amplicon sequences have been deposited in the National Center for Biotechnology Information (NCBI) database under the accession number PRJNA1461841, while the complete filtered ASV table, including ASV read counts and corresponding relative abundances for each sample, is provided in the Supplementary Material.

## 3. Results

### 3.1. Overview of reactor operation

A lab-scale CSTR was operated under anaerobic conditions for 297 days at 30 ± 1 °C, pH 5.5 ± 0.1, and a hydraulic retention time of 4 days, using lactate as the electron donor and energy source, and propionate as the electron acceptor. Over the course of the experiment, the lactate:propionate ratio was gradually adjusted from 1.2 to 4.8 mol/mol by changing the lactate or propionate concentrations. As a result, the total substrate concentration ranged from 200 to 530 mM. An overview of the reactor performance can be found in Figure 1 and Table 2. In all tested conditions, steady-state operation was reached within 20-30 days. In the liquid phase, valerate was the main odd-chain carboxylate with concentrations ranging from 36 to 107 mM, while heptanoate was only detected at concentrations below 6 mM. Similarly, caproate was the main even-chain carboxylate followed by butyrate and acetate, and no octanoate was observed at any point. Carbon and electron balances closed well (88 ± 7 % and 97 ± 8 %, respectively), indicating that all major fermentation products had been identified. In all cases, the majority of the incoming carbon ended up in the liquid products, since carbon dioxide and biomass represented only 6 ± 2 % and 8 ± 4 % of the carbon balance, respectively. Despite the low gas yields, off-gas composition remained stable over the whole experimental period, with average H_2_ and CO_2_ of 34 ± 2% and 64 ± 3%, respectively.

**Figure 1.**
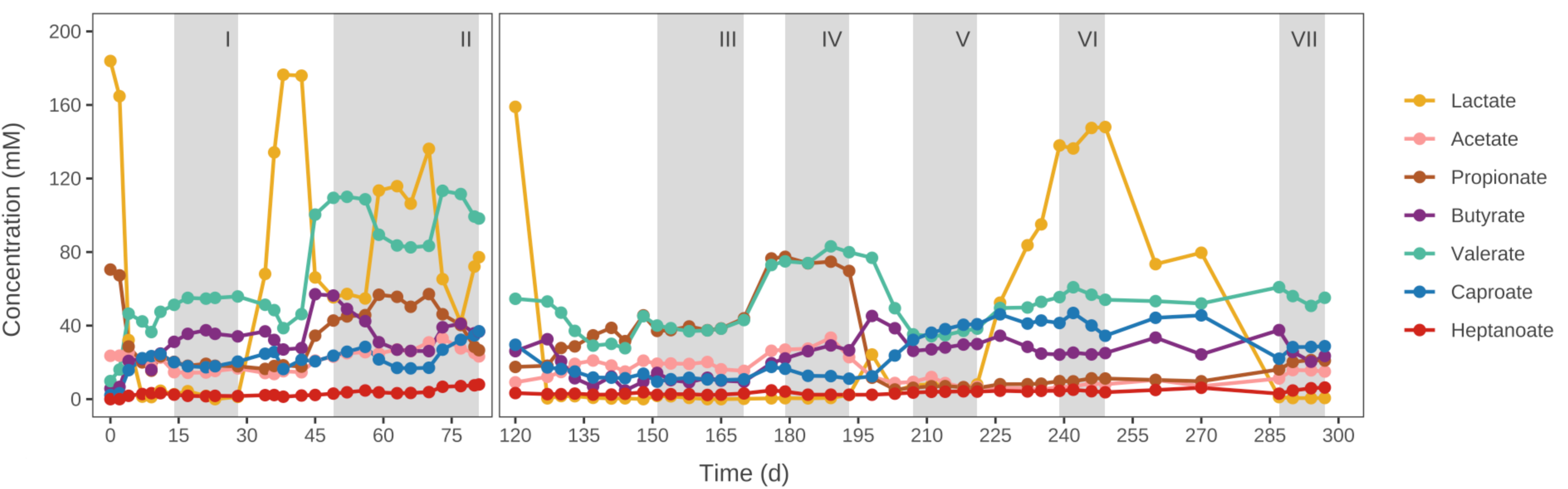
Overview of the reactor’s operation at different lactate:propionate molar ratios. Grey colour stands for steady state conditions from which the data was taken for comparison. Between day 81 and 120, there were operational issues and the data is not plotted.

**Table 2.** Carbon balance, product yields, and off-gas composition at the different conditions tested. Values represent the average and standard deviation of measurements taken during each steady-state period. “Others” represent isobutyrate, isovalerate and isocaproate.

| PHASE |  | I | II | III | IV | V | VI | VII |
| --- | --- | --- | --- | --- | --- | --- | --- | --- |
| Lactate:propionate ratio |  | 2.4 | 4.8 | 1.2 | 1.2 | 4.8 | 4.8 | 2.4 |
| Lactate in (mM) |  | 220 | 440 | 110 | 220 | 220 | 330 | 220 |
| Propionate in (mM) |  | 90 | 90 | 90 | 180 | 45 | 70 | 90 |
| Total concentration (mM) |  | 310 | 530 | 200 | 400 | 265 | 400 | 310 |
| Electron balance (%) |  | 89 ± 2* | 101 ± 3 | 103 ± 3 | 86 ± 5 | 105 ± 6 | 97 ± 3* | 95 ± 3* |
| Carbon balance (%) |  | 75 ± 2* | 88 ± 4 | 95 ± 4 | 81 ± 2 | 89 ± 4 | 78 ± 3* | 79 ± 3* |
| Product yield<br>(cmol/cmol) | Acetate | 0.03 ± 0.00 | 0.04 ± 0.00 | 0.08 ± 0.01 | 0.06 ± 0.01 | 0.02 ± 0.01 | 0.0 ± 0.00 | 0.04 ± 0.00 |
|  | Butyrate | 0.16 ± 0.01 | 0.13 ± 0.02 | 0.09 ± 0.01 | 0.11 ± 0.01 | 0.15 ± 0.01 | 0.10 ± 0.00 | 0.11 ± 0.01 |
|  | Valerate | 0.31 ± 0.02 | 0.41 ± 0.02 | 0.40 ± 0.02 | 0.39 ± 0.02 | 0.24 ± 0.01 | 0.30 ± 0.01 | 0.31 ± 0.01 |
|  | Caproate | 0.13 ± 0.01 | 0.14 ± 0.02 | 0.13 ± 0.01 | 0.08 ± 0.00 | 0.30 ± 0.02 | 0.26 ± 0.03 | 0.20 ± 0.00 |
|  | Heptanoate | 0.01 ± 0.00 | 0.03 ± 0.01 | 0.04 ± 0.00 | 0.02 ± 0.00 | 0.04 ± 0.00 | 0.03 ± 0.00 | 0.05 ± 0.01 |
|  | Others | 0.03 ± 0.00 | 0.02 ± 0.00 | 0.03 ± 0.01 | 0.02 ± 0.00 | 0.03 ± 0.00 | 0.03 ± 0.00 | 0.03 ± 0.00 |
|  | CO<sub>2</sub> | n.d. | 0.07 ± 0.01 | 0.05 ± 0.01 | 0.06 ± 0.01 | 0.07 ± 0.01 | n.d. | n.d. |
|  | Biomass | 0.08 ± 0.01 | 0.05 ± 0.01 | 0.14 ± 0.03 | 0.07 ± 0.01 | 0.09 ± 0.02 | 0.07 ± 0.01 | 0.08 ± 0.01 |
| Off-gas composition (%) | H<sub>2</sub> | 36 ± 1 | 36 ± 2 | 33 ± 1 | 33 ± 1 | 34 ± 1 | 31 ± 0 | n.d. |
|  | CO<sub>2</sub> | 60 ± 2 | 60 ± 1 | 61 ± 1 | 67 ± 0 | 67 ± 0 | 69 ± 1 | n.d. |

\* These carbon balances do not include the corresponding H<sub>2</sub> or CO<sub>2</sub>, as production rate was not determined.

### 3.2. Effect of lactate:propionate ratios on product spectrum

#### 3.2.1. At fixed lactate concentrations

The effect of the lactate:propionate molar ratio was evaluated in conditions with constant lactate concentration at 220 mM and varying propionate concentration from 180 mM (Phase IV) to 45 mM (Phase V) and 90 mM (Phase VII), as shown in Figure 1. Increasing propionate concentrations and thereby reducing the lactate:propionate ratio from 4.8 to 1.2 mol/mol, resulted in a higher fraction of odd-chain carboxylates compared to those produced at higher lactate:propionate ratios (Figure 2). At a molar ratio of 1.2, valerate was the dominant carboxylate, with selectivities up to 54 ± 3 %, followed by caproate and butyrate. Heptanoate was also detected under these conditions, albeit at concentrations of 2.8 ± 0.7 mM. On the contrary, at higher lactate:propionate molar ratios, the trend reversed, and even-chain carboxylates became the predominant product, reaching selectivities up to 65 ± 3 %. Valerate and heptanoate selectivities consequently decreased to only 35 ± 2 %.

**Figure 2.**
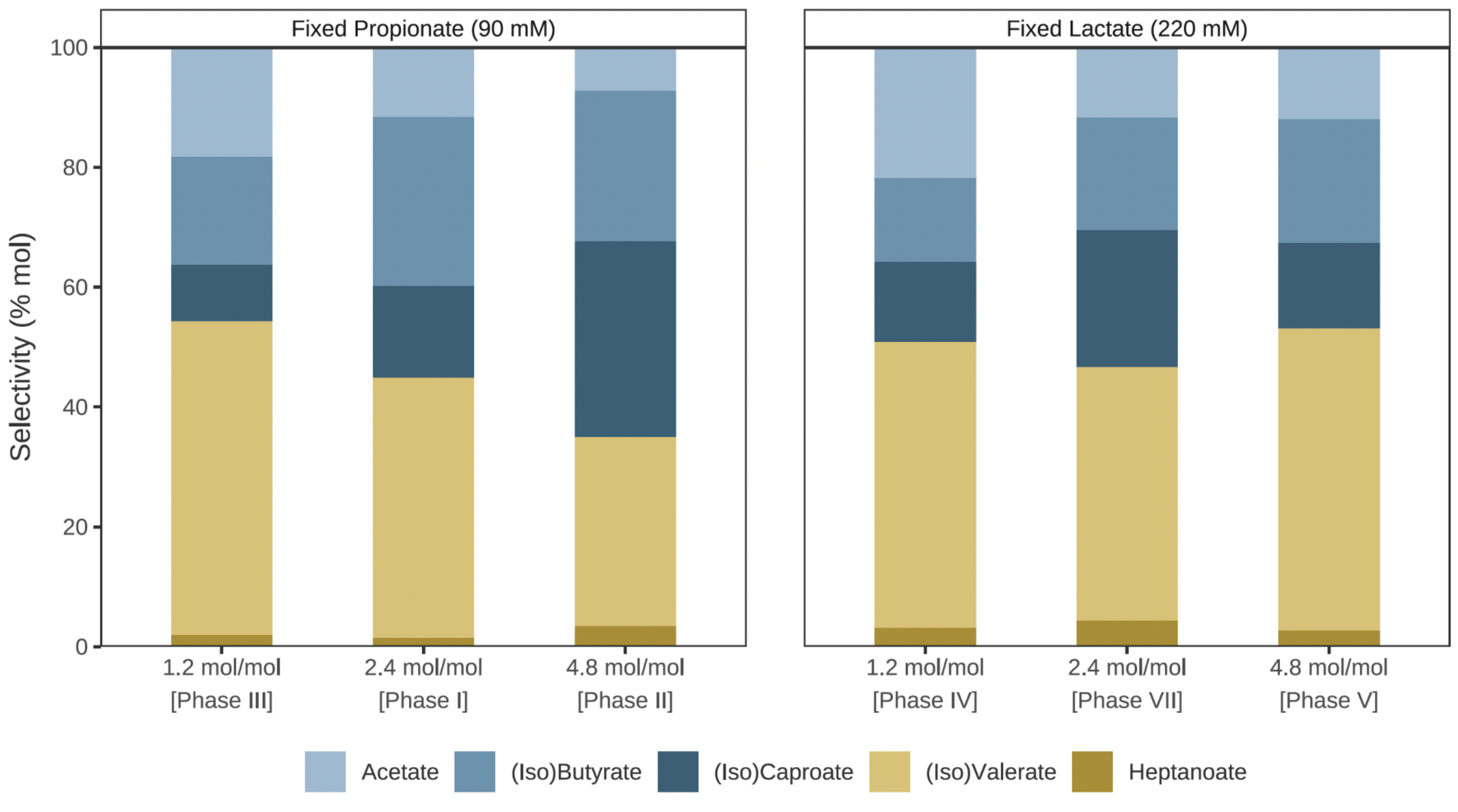
Product selectivities at varying lactate:propionate molar ratios and fixed lactate (220 mM) or propionate (90 mM) concentrations.

Yields on substrate followed similar trends: valerate and heptanoate yields increased from 0.24 ± 0.01 to 0.39 ± 0.02 cmol/cmol as the lactate:propionate ratio decreased, whereas caproate and butyrate yields increased at higher lactate:propionate ratios from 0.19 ± 0.01 to 0.45 ± 0.02 cmol/cmol, as illustrated in Table 2. Pathway analysis indicated that chain elongation was dominant throughout the tested conditions, while inferred endogenous propionate production was largely negligible, with a residual 1 ± 1% of the incoming lactate converted to acetate and propionate.

#### 3.2.2. At fixed propionate concentrations

The effect of the lactate:propionate molar ratio was further evaluated by comparing conditions with constant propionate concentration at 90 mM and varying lactate concentration from 110 mM (Phase III) to 220 mM (Phase I) and 440 mM (Phase II). This corresponded to an increase in the lactate:propionate molar ratio from 1.2 to 4.8 mol/mol (Figure 2). In contrast to the trends observed when varying propionate concentrations at fixed lactate levels, increasing the lactate concentration did not lead to a marked shift in product selectivity. Odd-chain selectivities at low and high lactate:propionate molar ratios remained comparable (51 ± 3 % and 53 ± 4 %, respectively), with the highest valerate concentrations (107 ± 6 mM) obtained under these conditions. Additionally, at a molar ratio of 4.8, up to 14 ± 2 % of the supplied lactate remained unconsumed, and its residual concentration fluctuated over time.

Yields on substrate also followed a similar pattern (Table 2). Valerate yields were comparable across the tested lactate:propionate molar ratios (0.41 ± 0.02 and 0.40 ± 0.02 cmol/cmol), while caproate and butyrate yields also showed limited variation (0.21 ± 0.01 and 0.27 ± 0.02 cmol/cmol). Pathway analysis indicated that, unlike in the previous experiment, the endogenous propionate production became significantly more active at high lactate:propionate ratios, with up to 24 ± 5 % of the incoming lactate diverted toward acetate and propionate formation, as shown in Figure 3.

**Figure 3.**
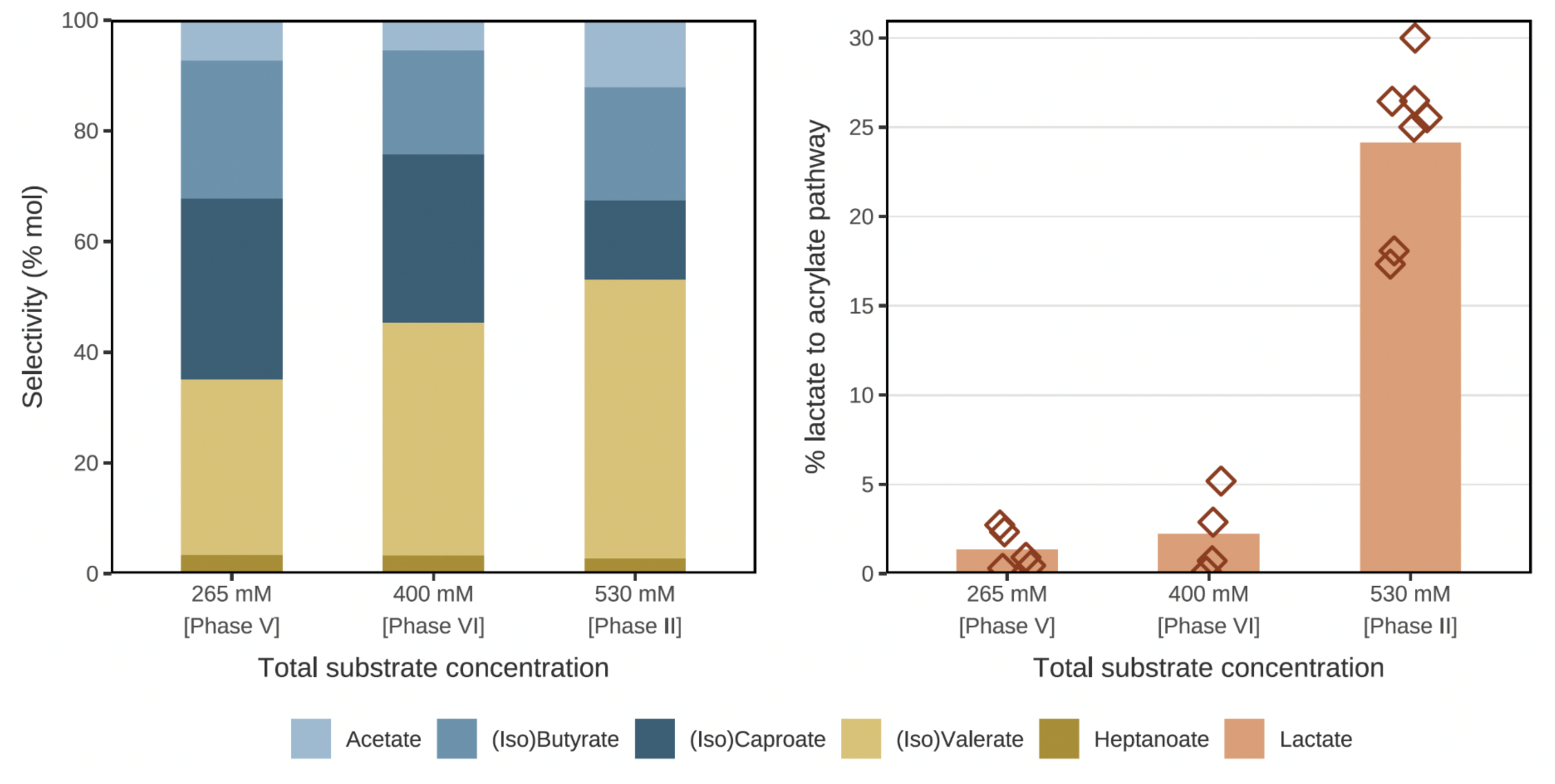
(A) Product selectivities at fixed lactate:propionate molar ratios of 4.8 but increasing total substrate concentrations; (B) Percentage of lactate used in inferred endogenous propionate production at each of the conditions described in (A). The inferred endogenous propionate production was calculated according to Equation 1.

### 3.3. Effect of lactate concentration on product spectrum at high lactate:propionate ratios

Because of the unexpectedly high valerate selectivities observed at high lactate:propionate molar ratios, it was hypothesized that the lactate concentration and/or lactate excess might influence endogenous propionate production. To further explore this theory, an additional condition (Phase VI) was tested at a lactate concentration of 330 mM, intermediate between 220 and 440 mM, while maintaining a constant lactate:propionate ratio of 4.8 mol/mol, as shown in Figure 3. Under these conditions, up to 43 ± 2 % of the supplied lactate remained unconsumed. Odd-chain selectivities decreased compared to the highest lactate concentration from 53 ± 4 to 44 ± 2 %, whereas even-chain selectivities increased from 47 ± 3 to 56 ± 3 %. Yields on substrate also followed similar patterns, but the endogenous propionate production remained negligible, and was only 2 ± 2 % of the incoming lactate. Overall, at a constant lactate:propionate molar ratio of 4.8, the fraction of odd-chain products increased with increasing lactate concentration, but not necessarily due to lactate excess.

### 3.4. Microbial community composition

Microbial community composition differed depending on whether the lactate:propionate ratio was altered by changing propionate or lactate concentrations, as observed in Figure 4. When the lactate:propionate ratio was modified by varying the propionate concentration and maintaining lactate constant at 220 mM, the community remained relatively stable at low and intermediate ratios, with *Caproiciproducens* spp. as the dominant genus (63–69% relative abundance). However, at higher lactate:propionate ratios, a pronounced community shift was observed. The relative abundance of *Caproiciproducens* spp. decreased to approximately 28%, coinciding with increased community diversity and a broader product spectrum. Genera such as *Dysgonomonas*, *Ignatzschineria*, and *Clostridium sensu stricto* increased to relative abundances above 10%. These changes coincided with the change from a product spectrum dominated by valerate to another dominated by butyrate and caproate. At the ASV level, *Caproiciproducens* ASV1 and ASV2 were the main ASVs and coexisted at lower and intermediate ratios, whereas only ASV1 persisted at higher ratios. BLAST analysis performed against the NCBI nucleotide database indicated that ASV2 showed 100% sequence identity (over the aligned region) to *Caproicibacterium lactatifermentans*, while ASV1 showed highest similarity to uncultured sequences affiliated with chain-elongating taxa, preventing further taxonomic resolution.

**Figure 4.**
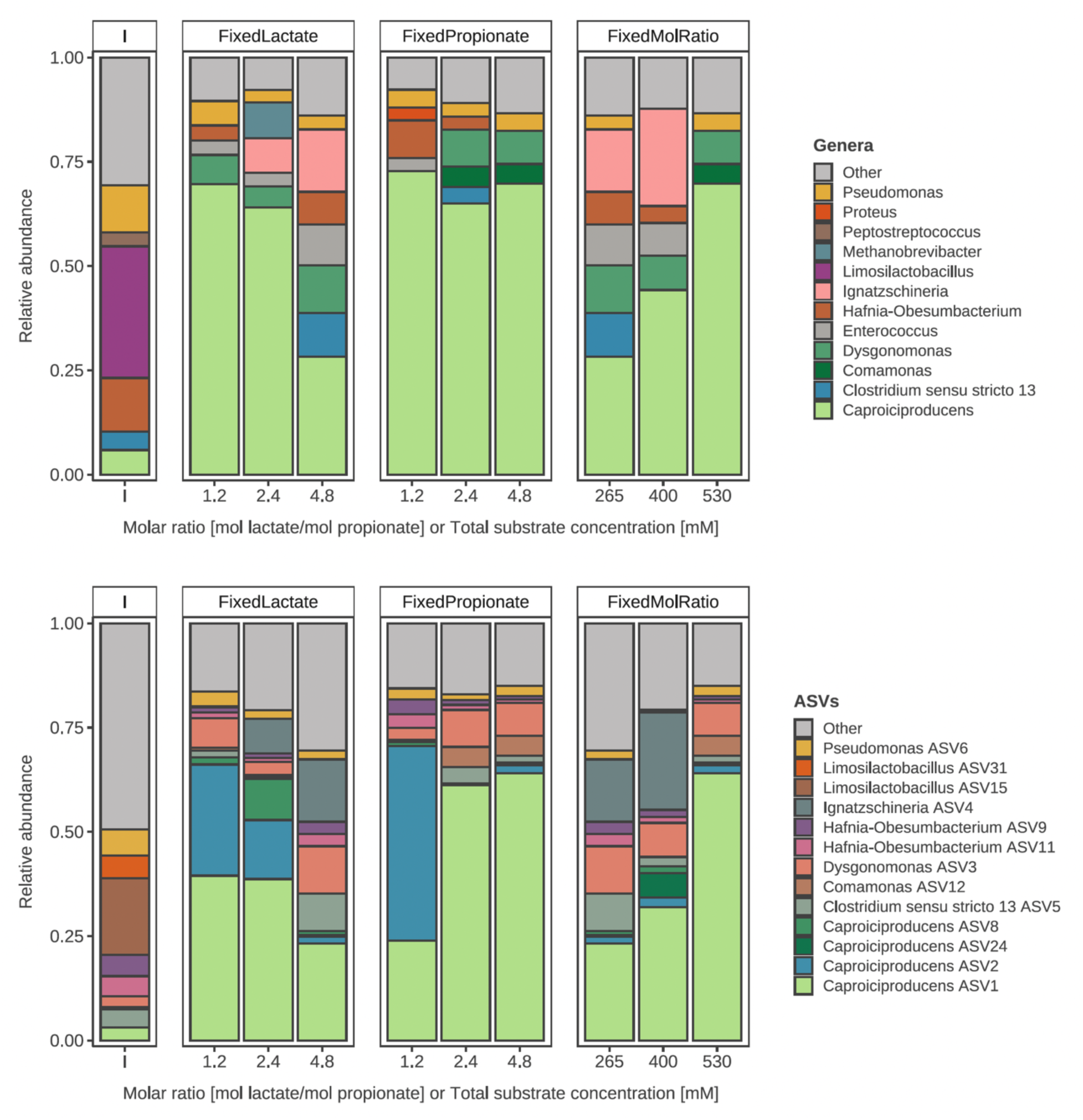
Microbial community structure under the different lactate:propionate molar ratios. On the top, the main genera are described, while at the bottom graph, the main ASVs are described. In both cases, relative abundances below 3% were lumped as Other.

In contrast, when the lactate:propionate ratio was adjusted by varying the lactate concentration and maintaining propionate constant, the microbial community remained comparatively stable across all tested ratios. *Caproiciproducens* spp. remained dominant, with relative abundances between 64 and 72%, and most other genera remained below 5%. As in the previous experiment, ASV1 and ASV2 coexisted at low ratios, while ASV1 dominated at intermediate and higher ratios. Under these conditions, valerate selectivities also remained comparable (51 ± 3 % and 53 ± 4 %). To further evaluate the effect of the lactate concentration, additional experiments were conducted at a fixed high lactate:propionate ratio (4.8 mol/mol) and higher lactate and propionate concentrations. Increasing lactate concentration resulted in a marked increase in the relative abundance of *Caproiciproducens* spp., from 28% to 68%, with ASV1 as the dominant ASV under these conditions.

## 4. Discussion

### 4.1. Valerate is the main product in lactate fermentations when propionate is present

Valerate was the main product across all tested conditions, except at higher lactate:propionate molar ratios and low lactate concentrations, where caproate became dominant. Valerate concentrations obtained in the present study ranged between 35 and 115 mM, while heptanoate was only detected at concentrations up to 6 mM, in line with previous studies^23,28,29^. Octanoate was never observed. Taken together, the results indicate that under acidic conditions and in the presence of odd-chain electron acceptors like propionate, lactate is predominantly channeled towards valerate rather than heptanoate. This trend has also been observed in engineered and natural environments. Previous studies using lactate as the sole electron donor have reported valerate as the main carboxylate, even when propionate was not externally supplied^9,18^. These patterns have also been observed in ethanol and methanol driven systems, where valerate is similarly the main carboxylate produced^23,28^. Valerate has also been detected in natural environments such as the human gut. Taxa commonly found in these environments, including *Lachnospiraceae* spp. and *Megasphaera* spp., contribute to valerate formation during secondary fermentations in which lactate is produced by lactic acid bacteria from readily available carbohydrates, while even and odd short-chain carboxylates are produced by other fermentative bacteria^20–22^.

Until recently, there was no clear mechanistic explanation for why shorter chain elongation intermediates (i.e., butyrate and valerate) are preferentially formed over longer-chain products (i.e., caproate, heptanoate, and octanoate). A recent study by Gois and co-workers provides new insights into the metabolic and genetic basis underlying this phenomenon^30^. Caproate producers tend to rely more on acetate recycling and substrate level phosphorylation, resulting in longer carboxylates to balance the additional redox equivalents from lactate oxidation, whereas butyrate producers appear to favor acetate assimilation and ion motive force for ATP synthesis through the RNF complex, enabling faster growth rates. Based on this reasoning, low acetate-to-lactate ratios should promote longer-chain carboxylates, whereas high acetate-to-lactate ratios should favor shorter ones. In the present study, no correlation was observed between the lactate:propionate ratio and the valerate:heptanoate ratio, suggesting that further research is required to clarify this point.

Product toxicity may offer an alternative explanation^31–34^. In acidic environments (pH < 6), a large fraction of acids remains in their undissociated form, increasing passive diffusion across the cell membrane and thereby raising the energetic burden required to maintain intracellular pH. For example, higher heptanoate concentrations compared to the present study have been reported in systems operated at pH 7^23^. However, the pKa values of valerate and heptanoate are very similar (valerate ≈ 4.8; heptanoate ≈ 4.9), making it unlikely that acid dissociation alone explains their differing abundances. Hydrophobicity has also been proposed as a quantitative predictor of metabolite toxicity^32^. At higher hydrophobicity, metabolites become more membrane soluble, where they disrupt membrane integrity and alter cell morphology^31,34^. Because hydrophobicity increases with carboxylate chain length, toxicity rises sharply from valerate (5 carbons) to octanoate (8 carbons). This is consistent with experimental observations showing that 50 mM of octanoate completely inhibited *E. coli* growth, whereas 50 mM of valerate caused only a 40% reduction in maximum growth rate^34^. Such differences in membrane impact offer a plausible explanation for why longer odd-chain carboxylates, such as heptanoate, accumulate to a lesser extent under acidic conditions. In the present study, the product spectrum remained similar across all operational conditions and consistently comprised butyrate, valerate and caproate as main products. Changes were mainly observed in their relative abundances and were associated with the lactate ratio rather than with increasing product concentrations. Therefore, although product toxicity cannot be excluded, there was no clear indication of product inhibition within the concentration range tested.

### 4.2. Lactate:propionate ratio drives product selectivity

The present study clearly shows that at low lactate concentrations, the lactate:propionate molar ratio drives product selectivity. At lower lactate:propionate molar ratios (higher propionate availability), odd-chain selectivity increased and valerate and heptanoate dominated the product profile. On the contrary, at higher lactate:propionate molar ratios (higher lactate availability), even-chain selectivity increased and caproate and butyrate dominated. This is in line with previous experimental work with ethanol and methanol as electron donor^28,35^. This trend is also consistent with the reverse β-oxidation pathway, where acetyl-CoA (formed from lactate) is combined with a propionyl-CoA molecule during odd-chain elongation to eventually yield valerate or heptanoate. However, for every 5 molecules of acetyl-CoA used to produce valerate, one is further oxidised to acetate to generate energy for the cell. This acetate is then also available to be elongated to butyrate and caproate. Consequently, assuming this stoichiometry, the higher the lactate compared to propionate, the higher the fraction of even-chain carboxylates. Theoretically, a maximum selectivity of around 83% is expected when the molar ratio is around 1.2 (6/5) mol lactate per mol of propionate. In the present study, at 1.2 molar ratio the selectivity towards odd-chain carboxylates was only 51-54%. This divergence from theoretical values has also been observed in ethanol-based chain elongation, where at molar ratios of 1.2, selectivities were also significantly lower than the theoretically expected, around 70%^35^. Candry et al. (2020) proposed that this may be due to a variable stoichiometry but could not pinpoint whether there could be different preferences for different electron acceptors. They also disproved the hypothesis that an increase in hydrogen partial pressure might limit the anaerobic oxidation of ethanol. Similarly, in the present study no correlation was found between lactate:propionate molar ratios and hydrogen partial pressures.

The changes in product selectivities were also reflected in the microbial community composition. *Caproiciproducens* spp. abundance tended to increase with valerate selectivity (Spearman’s ρ = 0.68, p = 0.094, n = 7), although this correlation was not statistically significant. *Caproiciproducens* spp. dominated the microbial community when valerate was the main product, reaching a 63–69% relative abundance (Figure 4). Similarly, when valerate production diminished, the relative abundance of *Caproiciproducens* spp. decreased to approximately 28%. *Caproiciproducens* spp. are known lactate-based chain elongators^36^ and are frequently reported in continuous enrichment systems^1,9,10,12,14^. It is therefore unclear why their relative abundance decreased at higher lactate:propionate ratios, as they are also known to be able to produce caproate from mixtures of lactate and acetate. This change suggests that the enriched *Caproiciproducens* spp. may be specialized in the use of propionate as electron acceptor. A decrease in *Caproiciproducens* spp. was however not observed at higher lactate concentrations, where odd chain selectivities were similar at both low and high lactate:propionate molar ratios; 51 ± 3 % and 53 ± 4 %, respectively. This trend was also observed in the microbial community composition, where *Caproiciproducens* spp. remained dominant, with relative abundances between 64 and 72%. This was unexpected, as higher lactate availability is typically associated with the production of longer-chain products (i.e. heptanoate) or a higher proportion of even-chain carboxylates due to lactate oxidation to acetate. Since heptanoate is likely toxic even at the low concentrations observed in the present study (0.83–1.54 mM undissociated heptanoate at pH 5.5), a higher fraction of even-chain carboxylates was therefore expected; however, this was not observed. Overall, this stable community composition suggests that the higher odd-chain selectivities obtained at high lactate:propionate ratios are likely due to a shift in intracellular carbon fluxes rather than a change in the microbial community composition.

### 4.3. Propionate is produced at high lactate concentrations

Counterintuitively, at high lactate concentrations more odd-chain carboxylates were obtained than theoretically expected, based on the amount of propionate consumed. Given that both valerate and heptanoate production require propionate as an initial electron acceptor, propionate had to be endogenously produced in an alternative way. Lactate can be converted to propionate (and acetate) via the acrylate or methylmalonyl-CoA pathways, which has been reported to also occur in conditions similar to those of the present study^9,14^. Endogenous propionate production could potentially explain this increase in odd-chain production. The pathway analysis showed that endogenous propionate production was significant only at high lactate concentrations, whereas it remained marginal at lower and intermediate lactate concentrations where lactate was fully consumed. This behavior contrasts with the expectation that higher lactate availability should always promote a greater fraction of even-chain elongation products due to increased electron donor oxidation to acetate. Instead, under high lactate:propionate molar ratios and lactate concentrations, odd-chain selectivity remained unexpectedly high, and up to 30 % of the incoming lactate was diverted to propionate and acetate. This endogenous formation of propionate compensated for the limited propionate supplied in the influent and, as a result, minimized the expected shift towards even-chain carboxylate formation.

As the microbial community composition remained fairly similar, it seems likely that propionate production results from a shift in intracellular carbon fluxes rather than a change in the microbial community composition. Similar observations have been made both in pure and mixed cultures, where propionate formation occurred exclusively under conditions of lactate excess. In *Megasphaera elsdenii*, Prabhu et al. (2012) showed that propionate was produced in batch cultures with excess lactate, but not under carbon-limited steady-state conditions in which lactate was fully consumed and butyrate and caproate were formed^37^. Likewise, Kucek et al. (2016) reported that in a continuous mixed-culture system, propionate production increased only when residual lactate accumulated, while no propionate was detected when lactate remained below the detection limit^14^. Similarly, Candry et al. (2020) reported significant odd-chain production (up to 35% of the incoming lactate) in a CSTR operated at pH 5.5, in which lactate was never fully consumed^9^. This explanation is also consistent with current resource-allocation theories, which predict that pathway use reflects a balance between ATP yield and proteome cost^38^. In *Megasphaera* spp., reverse β-oxidation provides a higher ATP yield (≈0.5 mol ATP per mol lactate) but requires multiple conversion steps and a larger enzymatic investment. In contrast, the acrylate pathway yields less energy (≈0.33 mol ATP per mol lactate), but its enzyme cost per ATP flux is estimated to be 4–7 times lower^38^. As a result, the acrylate pathway can support higher fluxes when lactate accumulates transiently, providing a proteome-efficient alternative under conditions of high lactate availability. In the present study, endogenous propionate production may therefore provide a proteome efficient alternative under high lactate conditions. However, lactate accumulation alone may not fully explain pathway activation, since the intermediate lactate concentrations showed substantial residual lactate but negligible inferred propionate formation.

The implications of this metabolic shift are that the lactate:propionate molar ratio controls odd-chain elongation only at low lactate concentrations. When endogenous propionate production becomes significant, this relationship breaks down, and product selectivity reflects the combined influence of external and endogenous propionate pools. These findings also introduce an opportunity to rewire carbon fluxes in lactate fermentations by activating an alternative propionate production pathway. From a bioprocess perspective, this can be of interest for PHBV production, where a higher HV content is desirable^39^. However, this also means that manipulating the influent lactate:propionate ratio alone may be insufficient to steer product selectivity in systems where lactate is abundant, such as sequencing batch reactors, due to the activation of propionate production pathways. Controlling lactate accumulation may therefore be necessary to suppress endogenous propionate formation and preserve expected odd-versus even-chain elongation behavior. Overall, the inferred propionate production at high lactate concentrations provides a mechanistic explanation for the deviations from expected product selectivities at elevated lactate:propionate ratios.

## 5. Conclusions

This study shows that lactate:propionate molar ratios and the substrate concentrations shape product selectivity in secondary lactate fermentations. Lower lactate:propionate ratios and full lactate consumption consistently increased odd-chain formation, with valerate as the dominant product. At higher lactate concentrations, this relationship broke down due to inferred endogenous propionate production, which redirected up to 30 % of the incoming lactate to propionate and acetate. This endogenous formation of propionate minimized the expected shift towards even-chain carboxylates and maintained valerate selectivities at 51-54 %. Overall, this work provides mechanistic insights into the ecology of lactate fermentations and may help steer the product spectrum in lactate fermentations by controlling lactate:propionate ratios and substrate concentrations.

## Data availability

Raw fastq files were deposited in NCBI under accession PRJNA1461841. All other data are included in this article and its supplementary information.

## Acknowledgments

The authors thank Anvesh Ganta for help setting-up the bioreactor.

## Author contributions

**Á. Estévez**: Conceptualization, Methodology, Investigation, Formal analysis, Visualization, Writing-Original Draft, Funding acquisition.

**R. Ganigué**: Conceptualization, Methodology, Writing-Review & Editing, Supervision, Funding acquisition.

## References

(1) Candry, P.; Ganigué, R. Chain Elongators, Friends, and Foes. Curr. Opin. Biotechnol. 2021, 67, 99–110. 10.1016/j.copbio.2021.01.005

(2) Agler, M. T.; Wrenn, B. A.; Zinder, S. H.; Angenent, L. T. Waste to Bioproduct Conversion with Undefined Mixed Cultures: The Carboxylate Platform. Trends Biotechnol. 2011, 29 (2), 70–78. 10.1016/j.tibtech.2010.11.006

(3) Angenent, L. T.; Richter, H.; Buckel, W.; Spirito, C. M.; Steinbusch, K. J. J.; Plugge, C. M.; Strik, D. P. B. T. B.; Grootscholten, T. I. M.; Buisman, C. J. N.; Hamelers, H. V. M. Chain Elongation with Reactor Microbiomes: Open-Culture Biotechnology To Produce Biochemicals. Environ. Sci. Technol. 2016, 50 (6), 2796–2810. 10.1021/acs.est.5b04847

(4) Wang, J.; Yin, Y. Biological Production of Medium-Chain Carboxylates through Chain Elongation: An Overview. Biotechnol. Adv. 2022, 55, 107882. 10.1016/j.biotechadv.2021.107882

(5) Louis, P.; Duncan, S. H.; Sheridan, P. O.; Walker, A. W.; Flint, H. J. Microbial Lactate Utilisation and the Stability of the Gut Microbiome. Gut Microbiome 2022, 3, e3. 10.1017/gmb.2022.3

(6) Pipyn, P.; Verstraete, W. Lactate and Ethanol as Intermediates in Two-phase Anaerobic Digestion. Biotechnol. Bioeng. 1981, 23 (5), 1145–1154. 10.1002/bit.260230521

(7) Spormann, A. M. Fermentative Metabolism. In Principles of Microbial Metabolism and Metabolic Ecology; Springer International Publishing: Cham, 2023; pp 215–269. 10.1007/978-3-031-28218-8_9

(8) Rombouts, J. L.; Kranendonk, E. M. M.; Regueira, A.; Weissbrodt, D. G.; Kleerebezem, R.; van Loosdrecht, M. C. M. Selecting for Lactic Acid Producing and Utilising Bacteria in Anaerobic Enrichment Cultures. Biotechnol. Bioeng. 2020, 117 (5), 1281–1293. 10.1002/bit.27301

(9) Candry, P.; Radić, L.; Favere, J.; Carvajal-Arroyo, J. M.; Rabaey, K.; Ganigué, R. Mildly Acidic pH Selects for Chain Elongation to Caproic Acid over Alternative Pathways during Lactic Acid Fermentation. Water Res. 2020, 186, 116396. 10.1016/j.watres.2020.116396

(10) Duber, A.; Zagrodnik, R.; Chwialkowska, J.; Juzwa, W.; Oleskowicz-Popiel, P. Evaluation of the Feed Composition for an Effective Medium Chain Carboxylic Acid Production in an Open Culture Fermentation. Sci. Total Environ. 2020, 728, 138814. 10.1016/j.scitotenv.2020.138814

(11) Brodowski, F.; Łężyk, M.; Gutowska, N.; Kabasakal, T.; Oleskowicz-Popiel, P. Influence of Lactate to Acetate Ratio on Biological Production of Medium Chain Carboxylates via Open Culture Fermentation. Sci. Total Environ. 2022, 851, 158171. 10.1016/j.scitotenv.2022.158171

(12) Carvajal-Arroyo, J. M.; Candry, P.; Andersen, S. J.; Props, R.; Seviour, T.; Ganigué, R.; Rabaey, K. Granular Fermentation Enables High Rate Caproic Acid Production from Solid-Free Thin Stillage. Green Chem. 2019, 21 (6), 1330–1339. 10.1039/C8GC03648A

(13) Contreras-Dávila, C. A.; Carrión, V. J.; Vonk, V. R.; Buisman, C. N. J.; Strik, D. P. B. T. B. Consecutive Lactate Formation and Chain Elongation to Reduce Exogenous Chemicals Input in Repeated-Batch Food Waste Fermentation. Water Res. 2020, 169, 115215. 10.1016/j.watres.2019.115215

(14) Kucek, L. A.; Nguyen, M.; Angenent, L. T. Conversion of L-Lactate into n-Caproate by a Continuously Fed Reactor Microbiome. Water Res. 2016, 93, 163–171. 10.1016/j.watres.2016.02.018

(15) Mariën, Q.; Ulčar, B.; Verleyen, J.; Vanthuyne, B.; Ganigué, R. High-Rate Conversion of Lactic Acid-Rich Streams to Caproic Acid in a Fermentative Granular System. Bioresour. Technol. 2022, 355, 127250. 10.1016/j.biortech.2022.127250

(16) Brodowski, F.; Łężyk, M.; Gutowska, N.; Oleskowicz-Popiel, P. Effect of External Acetate on Lactate-Based Carboxylate Platform: Shifted Lactate Overloading Limit and Hydrogen Co-Production. Sci. Total Environ. 2022, 802, 149885. 10.1016/j.scitotenv.2021.149885

(17) Allegue, T.; Rafay, R.; Chandran, S.; Amin, S. A.; Fowler, S. J.; Rodríguez, J. Lactate Addition Boosts Valerate Yields in Granular Mixed Culture Carbohydrate Fermentation. J. Environ. Chem. Eng. 2022, 10 (6), 108869. 10.1016/j.jece.2022.108869

(18) Rafay, R.; Allegue, T.; Fowler, S. J.; Rodríguez, J. Exploring the Limits of Carbohydrate Conversion and Product Formation in Open Mixed Culture Fermentation. J. Environ. Chem. Eng. 2022, 10 (3), 107513. 10.1016/j.jece.2022.107513

(19) Estévez-Alonso, Á.; Pei, R.; Van Loosdrecht, M. C. M.; Kleerebezem, R.; Werker, A. Scaling-up Microbial Community-Based Polyhydroxyalkanoate Production: Status and Challenges. Bioresour. Technol. 2021, 327, 124790. 10.1016/j.biortech.2021.124790

(20) McDonald, J. A. K.; Mullish, B. H.; Pechlivanis, A.; Liu, Z.; Brignardello, J.; Kao, D.; Holmes, E.; Li, J. V.; Clarke, T. B.; Thursz, M. R.; Marchesi, J. R. Inhibiting Growth of Clostridioides Difficile by Restoring Valerate, Produced by the Intestinal Microbiota. Gastroenterology 2018, 155 (5), 1495–1507.e15. 10.1053/j.gastro.2018.07.014

(21) Fitzgerald, B. G.; Govind, M.; Firth, I. J.; Herold, L.; Froment, J.; Vancuren, S. J.; Allen-Vercoe, E.; Kimber, M. S.; Sorbara, M. T. Members of Lachnospiraceae Produce Valerate and Caproate in Response to Short-Chain Fatty Acids. Microbiome 2025, 13 (1), 201. 10.1186/s40168-025-02220-9

(22) Huertas-Díaz, L.; Gezer, M. E.; Marietou, A.; Hosek, J.; Thams, L.; Dalgaard, L. B.; Hansen, M.; Schwab, C. Megasphaera Contributes to Lactate-Driven Valerate Production in the Human Gut. Microbiome 2025, 13 (1), 210. 10.1186/s40168-025-02207-6

(23) Grootscholten, T. I. M.; Steinbusch, K. J. J.; Hamelers, H. V. M.; Buisman, C. J. N. High Rate Heptanoate Production from Propionate and Ethanol Using Chain Elongation. Bioresour. Technol. 2013, 136, 715–718. 10.1016/j.biortech.2013.02.085

(24) Roels, J. A. Application of Macroscopic Principles to Microbial Metabolism. Biotechnol. Bioeng. 1980, 22 (12), 2457–2514. 10.1002/bit.260221202

(25) Klindworth, A.; Pruesse, E.; Schweer, T.; Peplies, J.; Quast, C.; Horn, M.; Glöckner, F. O. Evaluation of General 16S Ribosomal RNA Gene PCR Primers for Classical and Next-Generation Sequencing-Based Diversity Studies. Nucleic Acids Res. 2013, 41 (1), e1–e1. 10.1093/nar/gks808

(26) Callahan, B. J.; McMurdie, P. J.; Rosen, M. J.; Han, A. W.; Johnson, A. J. A.; Holmes, S. P. DADA2: High-Resolution Sample Inference from Illumina Amplicon Data. Nat. Methods 2016, 13 (7), 581–583. 10.1038/nmeth.3869

(27) Quast, C.; Pruesse, E.; Yilmaz, P.; Gerken, J.; Schweer, T.; Yarza, P.; Peplies, J.; Glöckner, F. O. The SILVA Ribosomal RNA Gene Database Project: Improved Data Processing and Web-Based Tools. Nucleic Acids Res. 2012, 41 (D1), D590–D596. 10.1093/nar/gks1219

(28) de Smit, S. M.; de Leeuw, K. D.; Buisman, C. J. N.; Strik, D. P. B. T. B. Continuous N-Valerate Formation from Propionate and Methanol in an Anaerobic Chain Elongation Open-Culture Bioreactor. Biotechnol. Biofuels 2019, 12 (1), 132. 10.1186/s13068-019-1468-x

(29) Veras, S. T. S.; Cavalcante, W. A.; Gehring, T. A.; Ribeiro, A. R.; Ferreira, T. J. T.; Kato, M. T.; Rojas-Ojeda, P.; Sanz-Martin, J. L.; Leitão, R. C. Anaerobic Production of Valeric Acid from Crude Glycerol via Chain Elongation. Int. J. Environ. Sci. Technol. 2020, 17 (3), 1847–1858. 10.1007/s13762-019-02562-6

(30) Gois, I. M.; Bowers, C. M.; Kim, B.-C.; Flick, R.; Lawson, C. E. Acetate Utilization Strategy in Chain-Elongating Bacteria Determines Butyrate versus Medium-Chain Carboxylate Production. Nat. Microbiol. 2026. 10.1038/s41564-026-02320-8

(31) Desbois, A. P.; Smith, V. J. Antibacterial Free Fatty Acids: Activities, Mechanisms of Action and Biotechnological Potential. Appl. Microbiol. Biotechnol. 2010, 85 (6), 1629–1642. 10.1007/s00253-009-2355-3

(32) Jarboe, L. R.; Royce, L. A.; Liu, P. Understanding Biocatalyst Inhibition by Carboxylic Acids. Front. Microbiol. 2013, 4. 10.3389/fmicb.2013.00272

(33) Vázquez, J.; Durán, A.; Rodríguez-Amado, I.; Prieto, M.; Rial, D.; Murado, M. Evaluation of Toxic Effects of Several Carboxylic Acids on Bacterial Growth by Toxicodynamic Modelling. Microb. Cell Factories 2011, 10 (1), 100. 10.1186/1475-2859-10-100

(34) Wilbanks, B.; Trinh, C. T. Comprehensive Characterization of Toxicity of Fermentative Metabolites on Microbial Growth. Biotechnol. Biofuels 2017, 10 (1), 262. 10.1186/s13068-017-0952-4

(35) Candry, P.; Ulcar, B.; Petrognani, C.; Rabaey, K.; Ganigué, R. Ethanol:Propionate Ratio Drives Product Selectivity in Odd-Chain Elongation with Clostridium Kluyveri and Mixed Communities. Bioresour. Technol. 2020, 313, 123651. 10.1016/j.biortech.2020.123651

(36) Joshi, S.; Robles, A.; Aguiar, S.; Delgado, A. G. The Occurrence and Ecology of Microbial Chain Elongation of Carboxylates in Soils. ISME J. 2021, 15 (7), 1907–1918. 10.1038/s41396-021-00893-2

(37) Prabhu, R.; Altman, E.; Eiteman, M. A. Lactate and Acrylate Metabolism by Megasphaera Elsdenii under Batch and Steady-State Conditions. Appl. Environ. Microbiol. 2012, 78 (24), 8564–8570. 10.1128/AEM.02443-12

(38) Strachan, C. R.; Bowers, C. M.; Kim, B.-C.; Movsesijan, T.; Neubauer, V.; Mueller, A. J.; Yu, X. A.; Pereira, F. C.; Nagl, V.; Faas, J.; Wagner, M.; Zebeli, Q.; Weimer, P. J.; Candry, P.; Polz, M. F.; Lawson, C. E.; Selberherr, E. Distinct Lactate Utilization Strategies Drive Niche Differentiation between Two Co-Existing *Megasphaera* Species in the Rumen Microbiome. ISME J. 2025, 19 (1), wraf147. 10.1093/ismejo/wraf147

(39) Pei, R.; De Vries, E.; Estévez, A.; Sousa, J.; Dijkman, H.; Tamis, J.; Werker, A. Demonstrating Performance in Scaled-up Production and Quality Control of Polyhydroxyalkanoates Using Municipal Waste Activated Sludge. Water Res. 2025, 275, 123160. 10.1016/j.watres.2025.123160

